# Influenza A(H1N1)pdm09 M and HA segments sequences from Rio Grande do Sul, Brazil

**DOI:** 10.1101/347229

**Authors:** F. Matias, N.A. Kretzmann, C.Z. Bisognin, K.P. Teixeira, L.T. Corrêa, P.I. Vieira, N. Ikuta, T.S. Gregianini, A.B.G da Veiga, P.A. d’Azevedo

**Author notes:** Corresponding Author: Ana Beatriz Gorini da Veiga -, Universidade Federal de Ciências da Saúde de Porto Alegre (UFCSPA), R. Sarmento Leite, 245 / sala 309-Centro, CEP 90.050-170, Porto Alegre, RS, Brazil. Fernanda Matias -, Universidade Federal Rural do Semi-Árido, Rua Francisco Mota, 572, Departamento de Biociências, sala 25, Costa e Silva, CEP 59.625-900, Mossoró, RN, Brazil. Numbers of the deposited sequences: KU143688–KU143706 (M segment); MG784976–MG784982; MG785397–MG785406 (HA segment).

## Abstract

The influenza virus is one of the most critical viruses in epidemiology. The 2009 pandemic was caused by a reassortment of the human–avian–swine virus with eight RNA segments responsible for all virus proteins. Segment 7 codifies for matrix proteins M1 and M2. These proteins exhibited low mutation rate because the matrix is fundamental for virion encapsidation and ion channel formation. However, hemagglutinin (HA) segment 4 is one of the most important segments for virulence and hence, is more studied. Brazil had many influenza virus infection cases just before 2009 and from 2011 to 2015, particularly in the Rio Grande do Sul (RS) State. Two hundred samples obtained during the pandemic were used for amplification and sequencing of the viral genome; a total of 19 M and 17 HA amplified segments were sequenced. Sequencing of the M fragment showed that RS has a virus origin different from that in Eastern Asia, Western Europe, USA, and Central America (Mexico and Nicaragua). All the sequences showed amantadine resistance (S31N) and one was out of the phylogenetic tree (Brazil/RS-3335/2009) due to high mutation rate. RS-3335 was the only sample obtained from a patient who died. Many migratory birds that flock to RS are from Europe, Asia, and USA, which could explain this rate of mutation. Insertions and deletions were found in the M1 protein in these samples. The HA sequences showed worldwide spread and less diversity than the M sequences in this study. The most divergent sample was Brazil/RS-3093/2009 that showed mutations in the sialic acid ligation site.

## Introduction

The influenza virus is an enveloped RNA virus surrounded by a lipid bilayer that belongs to the *Orthomyxoviridae* family. This virus is responsible for the most significant pandemics worldwide and is first classified into A, B, or C types, with the last two being less common. Influenza A is divided according to antibody response to the combination between hemagglutinin (HA) and neuraminidase (NA) proteins, called “H” and “N” for this classification, respectively. There are at least 16 serotypes of H and nine serotypes of N, enabling 144 combinations. The segments H1, H2, and H3 combined with N1 and N2 serotypes are more common in humans (Lynch and Walsh 2007). In 2013, a new influenza A virus (H7N9) was described (WHO 2013). Among these serotypes, H1N1 was responsible for the 1918 (Spanish flu; Mills *et al*. 2004) and 2009 (swine flu; Schnitzler and Schnitzler 2009) pandemics.

The 2009 pandemic H1N1 (pH1N1) virus was first described in March in the cities of Sao Luís do Potosi and Oaxaca, Mexico. On April 17, 2009, it reached the United States of America, thus, spreading quickly worldwide, resulting in approximately 9 000 deaths by November of that year (Smith *et al*. 2009). In Brazil, the first case was reported on April 24 in Sao Paulo. Almost at the same time, one more case was reported in Sao Paulo, one case in Rio de Janeiro, and one in Minas Gerais—all southeastern states— with three patients coming from Mexico and one from USA. The first incident of death occurred in RS, the southernmost state of the country, on July 26. The patient was a truck driver who was in Argentina for seven days. RS had the highest number of cases and deaths, particularly during autumn and winter periods.

H1N1 is a virus consisting of eight RNA segments encoding eleven genes and proteins, viz., RNA polymerase basic proteins (PB1, segment 1; PB1-F2; and PB2, segment 2), RNA polymerase (PA; segment 3), hemagglutinin (HA; segment 4), nucleocapsid protein (NP; segment 5), neuraminidase (NA; segment 6), matrix proteins (M1 and M2; segment 7), and nonstructural proteins (NS1 and NS2; segment 8) (Ghedin 2005). HA and NA are the most studied proteins in influenza resistance and virulence, as these proteins are involved in the host immune response and are more likely to mutate. Matrix protein (MP) is divided into two proteins: M1, which is inside the envelope and interacts with the genome, synthesizing new viral fragments; and M2, which synthesizes ionic channels that facilitate the interaction between the inner and outer environments of the virus, primarily when it is producing HA and during viral desencapsidation. Unlike HA and NA, MP typically remains conserved, as the matrix is an essential part of the virus.

One copy of each segment is packaged during viral assembly in one virion particle. The PB2, PA, NP, and M segments are the most critical segments in virus packaging (Gao *et al*. 2012). According to Gao *et al*., (2012), Amorim *et al*. (2011), and Marsh *et al*. (2008), PB1, HA, NA, and NS play non-essential roles during genome assembly and may be more permissive toward reassortment than PB2, PA, NP, and M. The matrix segment has been studied more since these discoveries were made. This study aimed to obtain more data on the 2009 pandemic in the state of RS, Brazil, through M and HA sequence analysis and analysis of the relationship between the symptoms. Brazilian epidemiological data were also used to understand some of our results.

## Materials and Methods

### Samples and sequencing

Nasopharyngeal aspirate samples were collected from RS, Brazil, during the 2009 outbreak (Veiga *et al*. 2012). Ethics approval was provided by the Research Ethics Committee of Universidade Federal de Ciências da Saúde de Porto Alegre (UFCSPA). Identification of pH1N1 was conducted at the Central Laboratory of the State (LACEN) by real-time PCR (Veiga *et al*. 2012). RNA extractions were performed at Universidade Luterana do Brasil (ULBRA), using NewGene Extraction Kit (Simbios Biotecnologia, RS, Brazil), and handled at Universidade Federal de Ciências da Saúde de Porto Alegre (UFCSPA). cDNA library was constructed using SuperScript III Reverse Transcriptase according to the manufacturer’s instructions (Life Technologies). The samples were stored at −80°C.

Many approaches were used to amplify the HA and M segments, as mentioned in Table 1. Conventional PCR was conducted using KAPA 2G Robust HotStart ReadyMix (Biosystems, Boston, USA). The amplicons were treated with Shrimp Alkaline Phosphatase and Exonuclease I, marked with Big Dye Terminator v3.1 kit Cycle Sequencing Kit (Life Technologies), and sequenced using ABI 3130 (Life Technologies). The sequences were constructed and analyzed using Staden 2.0 for Windows and Pregap4 software (Staden 1996). First, all the H1N1 sequences were collected from the Influenza Research Database (IRD) (http://www.fludb.org/), aligned using the online software MAFFT and the FF-NS-2 strategy (http://mafft.cbrc.jp/alignment/server/), and a rough tree was constructed. Sequences with maximum similarity were selected for a new alignment. Next, the resulting sequences from this alignment were compared using BLAST (Basic Local Alignment Tool). All matches obtained in this search were used for a new alignment, using the MEGA 7 software (Tamura et al., 2016) and MUSCLE.

**Table 1:**
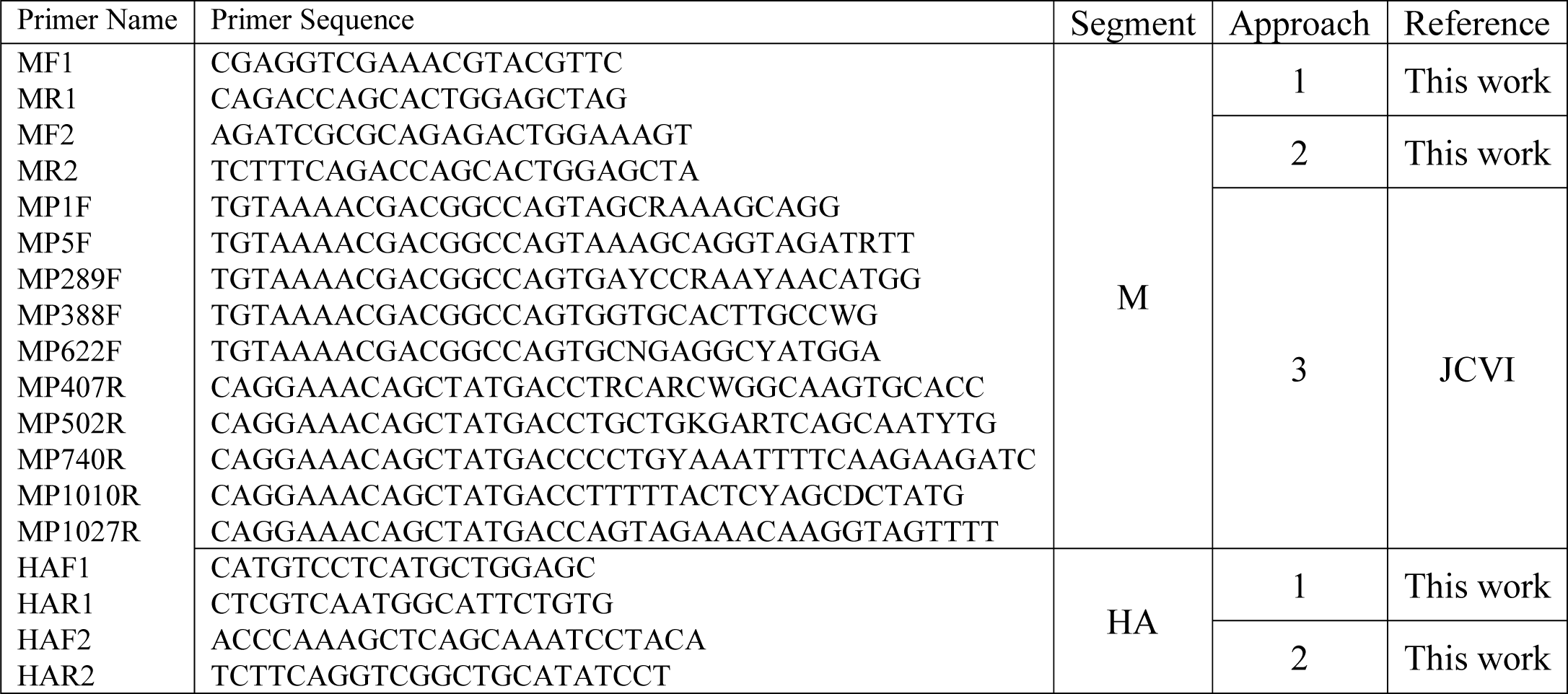
Primers used in this work.

### Phylogenetic analysis and map construction

The phylogenetic tree was constructed using the MEGA 7 software (Tamura *et al*. 2016) by the neighbor-joining (NJ) method (Saitou and Nei 1987) with the statistical method of maximum likelihood (ML) (Tamura *et al*. 2004). The trees were exported as Newick trees to the iTOL online software (Letunic and Bork 2011) to select subtrees to be used for new alignments. The shared phylogenetic trees can be found at the iTOL page, under the login ID fernandamatias. To construct an interactive map, all sequences were distributed in Excel according to their similarity and clade. This approach provided a better view of the sequences analyzed in this study. Information on GenBank accession number, host, strain, influenza type, and the location of samples were added to construct a more concise and complete map. The subtrees selected were used to create a map of RS samples and other worldwide more similar samples. The interactive map was constructed using the online platform Google My Maps (https://www.google.com/mymaps/).

### Epidemiological Influenza information

During the 2009 influenza pandemic, 218 pandemic influenza samples from RS (Veiga *et al*. 2011) were analyzed for acute respiratory infection (ARI) symptoms (fever, cough, chills, dyspnea, sore throat, arthralgia, myalgia, conjunctivitis, rhinorrhea, and diarrhea) and comorbidities (cardiopathy, pneumonia, renal complications, immunodepression, and chronic metabolic disease) using Fisher’s exact test to evaluate the relationships between the symptoms. p < 0.05 was considered to indicate statistically significant differences. The Gephi 0.9.1 software (Bastian *et al*. 2009) was used to analyze the most frequent and most related symptoms. Another approach was to analyze Brazilian online bulletins of influenza provided by the Brazilian Health Ministry (Brasil 2012a; Brasil 2012b; Brasil 2013; Brasil 2014; Brasil 2015) to understand the epidemiology of influenza in Brazil from 2009 to 2015. These bulletins were used to distinguish the number of cases and deaths from 2009 to 2015 in different regions of the country (Table 2).

**Table 2:**
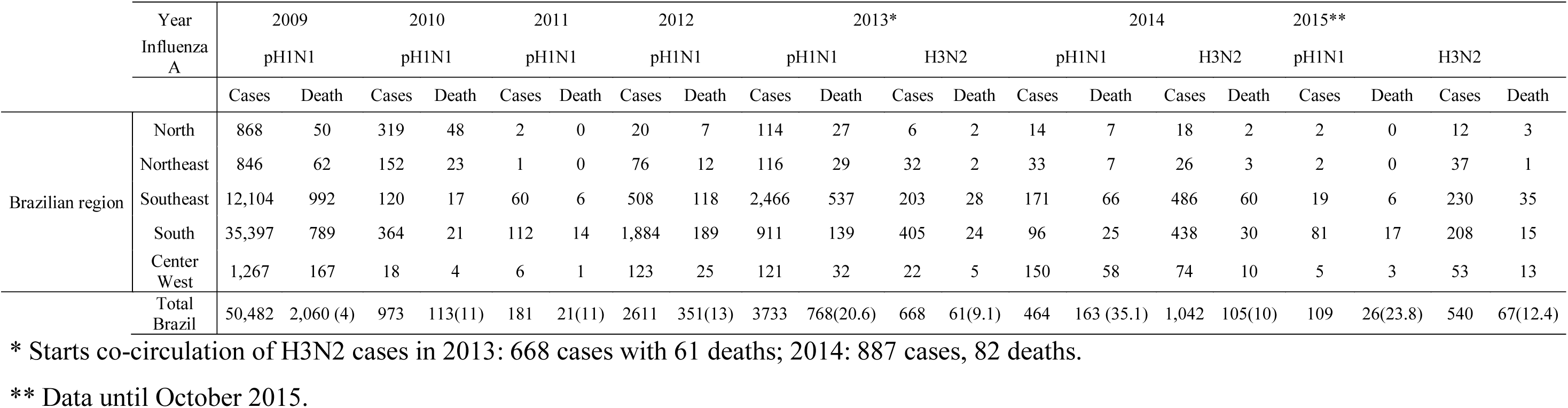
Number of cases and death caused by influenza A pH1N1 in Brazil from 2009 to 2015.

|  | Year<br>Influenza<br>A | 2009 |  | 2010 |  | 2011 |  | 2012 |  | 2013* |  |  |  | 2014 |  |  |  | 2015** |  |  |  |
| --- | --- | --- | --- | --- | --- | --- | --- | --- | --- | --- | --- | --- | --- | --- | --- | --- | --- | --- | --- | --- | --- |
|  |  | pH1N1 |  | pH1N1 |  | pH1N1 |  | pH1N1 |  | pH1N1 |  | H3N2 |  | pH1N1 |  | H3N2 |  | pH1N1 |  | H3N2 |  |
|  |  | Cases | Death | Cases | Death | Cases | Death | Cases | Death | Cases | Death | Cases | Death | Cases | Death | Cases | Death | Cases | Death | Cases | Death |
| Brazilian region | North | 868 | 50 | 319 | 48 | 2 | 0 | 20 | 7 | 114 | 27 | 6 | 2 | 14 | 7 | 18 | 2 | 2 | 0 | 12 | 3 |
|  | Northeast | 846 | 62 | 152 | 23 | 1 | 0 | 76 | 12 | 116 | 29 | 32 | 2 | 33 | 7 | 26 | 3 | 2 | 0 | 37 | 1 |
|  | Southeast | 12,104 | 992 | 120 | 17 | 60 | 6 | 508 | 118 | 2,466 | 537 | 203 | 28 | 171 | 66 | 486 | 60 | 19 | 6 | 230 | 35 |
|  | South | 35,397 | 789 | 364 | 21 | 112 | 14 | 1,884 | 189 | 911 | 139 | 405 | 24 | 96 | 25 | 438 | 30 | 81 | 17 | 208 | 15 |
|  | Center West | 1,267 | 167 | 18 | 4 | 6 | 1 | 123 | 25 | 121 | 32 | 22 | 5 | 150 | 58 | 74 | 10 | 5 | 3 | 53 | 13 |
|  | Total Brazil | 50,482 | 2,060 (4) | 973 | 113(11) | 181 | 21(11) | 2611 | 351(13) | 3733 | 768(20.6) | 668 | 61(9.1) | 464 | 163 (35.1) | 1,042 | 105(10) | 109 | 26(23.8) | 540 | 67(12.4) |

\* Starts co-circulation of H3N2 cases in 2013: 668 cases with 61 deaths; 2014: 887 cases, 82 deaths.
\*\* Data until October 2015.

## Results

### Genetic analysis

Several primers were tested for partial amplification of the M and HA segments, as shown in Table 1. First positive results for the M segments were obtained using MF1 and MR1 primers yielding a 700-bp fragment, but from all samples, only three segments were amplified. The second approach using MF2 and MR2 provided only seven amplifications of a 627-bp fragment, which was still not a good yield. Phylogenetic analysis of these amplifications showed a highlighted clade compared to that in samples available on genomics databases. Thus, it was necessary to obtain a new amplicon covering the largest possible area of the segment, as the highlighted clade could be an artifact of the amplified region with a small sample size. In this way, the third approach was designed to achieve complete M segment amplification. Of all the primers suggested by the John Craig Venter Institute (JCVI), 10 were synthesized and combined to yield nine pairs that were tested (Table 3). The combinations and number of amplified samples are summarized in Table 3. From all the M sequences obtained, only 19 sequences overlapped and were used to create a contiguous consensus sequence. For the HA sequence, the first approach resulted in five positives using HAF1 and HAR1, but the second approach using HAF2 and HAR2 yielded 17 amplicons of approximately 700 bp. Primers from JCVI were used as well but showed no positive amplification in these samples.

**Table 3:** Strategy of primers used in this work.

| Primers | Amplicon size | Strategy | Result |
| --- | --- | --- | --- |
| HAF1 + HAR1<br>MF1 + MR1 | ~700 bp<br>~700 bp | 1 | Few positives |
| HAF2 + HAR2<br>MF2 + MR2 | 627 bp<br>625 bp | 2 | Few positives, more than strategy 1 |
| MP1F + MP407R | 406 bp | 3 | Negative |
| MP1F + MP740R | 739 bp |  | Negative |
| MP5F + MP502R | 497 bp |  | Negative |
| MP289F + MP740R | 451 bp |  | Positive |
| MP388F + MP1010R | 633 bp |  | Positive |
| MP388F + MP1027R | 639 bp |  | 40 positives |
| MP622F + MP1027R | 405 bp |  | Positive |
| MF1 + MP740R | 739 bp |  | 30 positives |

Shadow: primers used to obtain partial M and HA segment.

The M sequences obtained from the RS samples received GenBank accession numbers KU143688–KU143706 and were compared to all sequences available in the genetic databases until October 2014. First, the RS sequences were aligned to the H1N1 sequences, human or not, and opened in IRD, irrespective of the year. Sequences that were more similar to the sequences obtained in this study were chosen and retrieved. Next, the RS sequences were analyzed one by one by BLAST to search for more similar sequences available in this database. With these two approaches, 466 sequences were obtained and compared to the RS sequences. All Brazilian sequences were included. The phylogenetic tree showed few branches (red circles) with the RS sequences (highlighted in yellow) (Figure 1). Two RS sequences were outgrouped in this analysis: Brazil/RS-3335/2009 and Brazil/RS-3504/2009. Seventy-four sequences, including 19 RS sequences, were selected for a new alignment and for generation of a new phylogenetic tree (Figure 2). The only significant difference in all sequences was obtained in sample Brazil/RS-3335/2009, which resulted in the death of the patient. For samples from Brazil, only one similarity was observed to three other RS sequences and one sequence from Sao Paulo, but only two of these sequences were related to the M sequences of this study. This phylogenetic tree was used to generate the interactive map to assess the most similar sequences associated with the result of this study (https://www.google.com/maps/d/edit?mid=zQfm5yljIn54.k7PW6v6h1pJI&usp=sharing). This map showed the position of the RS samples, almost all from the northeast of the state, with some being near Argentina (Figure 3). From all 60 Brazilian sequences, only two swine sequences (Figure 3; black square), Brazil/G2P2/2013 and Brazil/G2P1/2013, both H1N2, from RS were directly related to Brazil/RS-3538/2009 and Brazil/RS-3093/2009 sequences. Another significant result obtained was the global position of most similar sequences linked to the sequences of this study, which were from USA, Mexico, Western Europe, and Eastern Asia (Figure 4). Interactive map shows that there were no RS sequences against similar sequences from group 14. The most significant changes in amino acid composition were in M1 protein (Table 4). For the construction of M1 protein table, Brazil/RS-3335/2009 and Brazil/RS-3504/2009 sequences were excluded as they were vastly different from California/04/2009 and were more similar to Guangdong/55/2009. This approach was not necessary when comparing the M2 protein. The M1 protein showed insertions, deletions, and substitutions. Brazil/RS-1892/2009 and Brazil/RS-2009/2009 showed many similarities, including substitutions at G73A, E197T, and Q198T and insertion of R at position 74/75, K at position 197/198, and S at position 200/201. Brazil/RS-2656/2009 showed substitutions at A96T, S196R, and Q198T; deletion of E at position 197; and insertion of S at position 200/201. Brazil/RS-2905/2009 showed only one substitution, N91Q. Brazil/RS-3538/2009 showed two substitutions at the beginning of the protein, K47T and P54Y. M2 revealed more similarities than M1. All RS M sequences contained the signature mutation S31N in M2 that may have conferred amantadine resistance. Brazil/RS-3435/2009 showed the highest number of substitutions, viz., Q77N, D85G, V86C, and F91L. Brazil/RS-2014/2009, Brazil/RS-2656/2009, and Brazil/RS- 3335/2009 showed one amino acid substitution each, E95K, R54L, and I94K, respectively.

**Figure 1:**
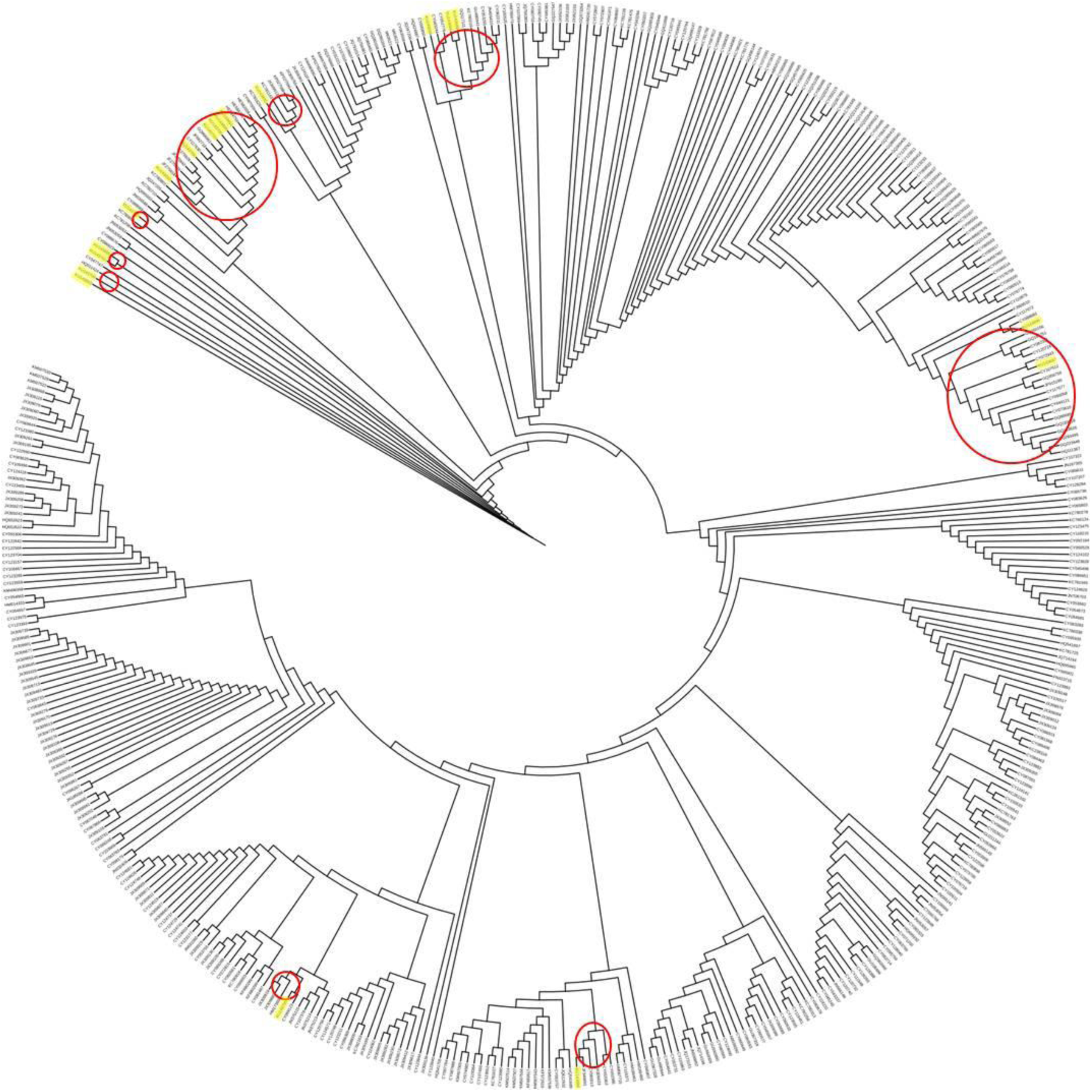
second selection with 466 sequences all over the world. All sequences of the branch with some of the RS sequences (yellow squares) were selected (red circles) to the new alignment.

**Figure 2:**
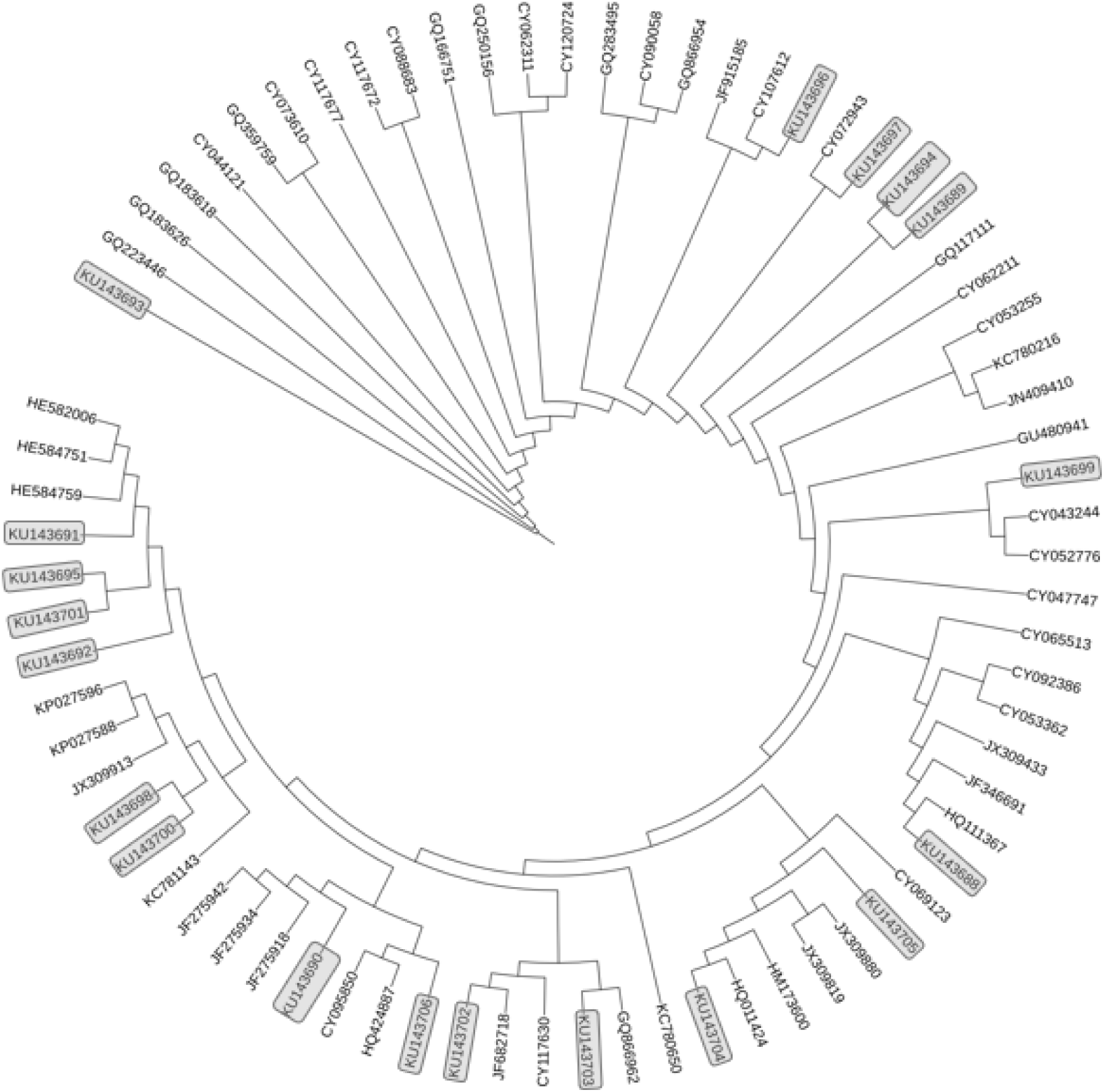
Final phylogenetic tree with 55 of the most similar sequences related to 19 RS sequences (gray squares). The most divergent sequence is KU143693 (Brazil/RS- 3335/2009) nearby sequences from Beijing.

**Figure 3:**
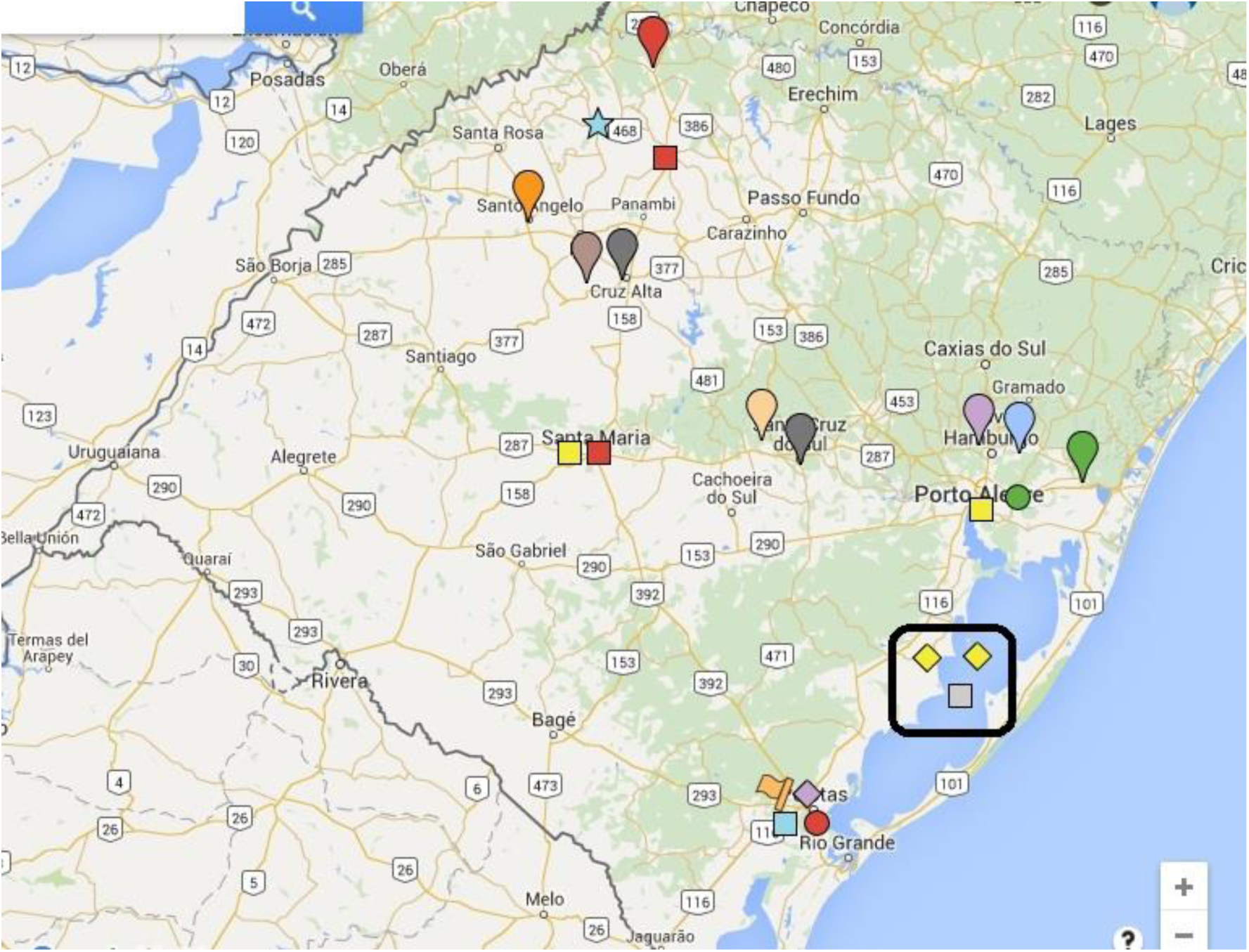
Rio Grande do Sul Map. It is interactive, click the link below and open image. Clicking icons, it is possible to see all information about data. The same color is the same branch. Drops are the first level, squares are the second level, circles are the third level, a rhombus is the fourth level, stars are the fifth level, and flags are the sixth level of the same branch of the tree. It is possible to see more similarity between samples analyzed in this study than from others. There are only three from RS sequences similar to those from theses work (Black Square), and the most similar are from swine and not human as yellow color shows. (https://www.google.com/maps/d/edit?mid=zQfm5yliIn54.k7PW6v6h1pJI&usp=sharing)

**Figure 4:**
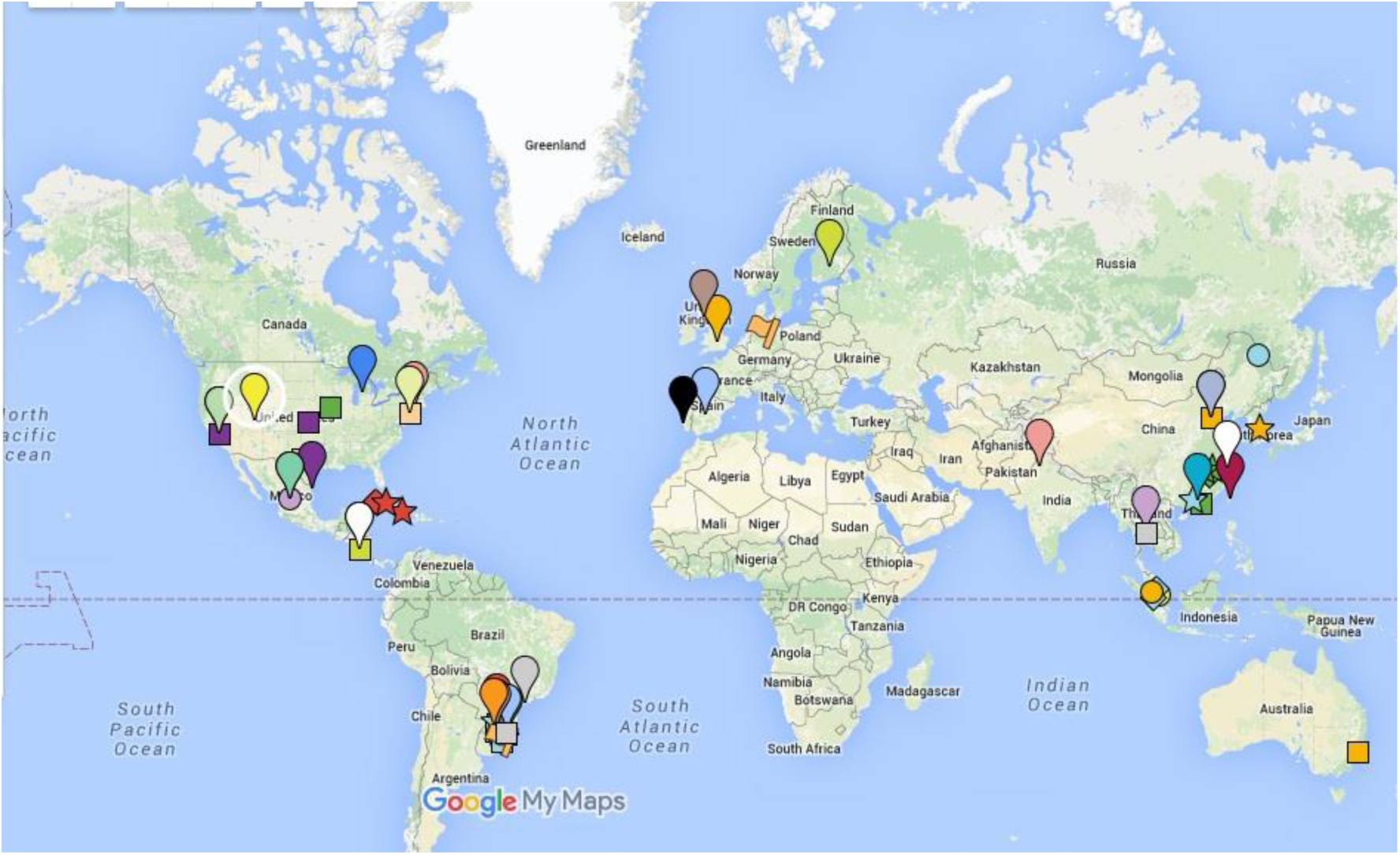
World Map constructed using phylogeny tree. It is interactive, click the link below and open image. Clicking on the icons, it is possible to see all information about data. The same color is the same branch. Drops are the first level, squares are the second level, circles are the third level, a rhombus is the fourth level, stars are the fifth level, and flags are the sixth level of the same branch of the tree. It is possible to see that most similarities were obtained to sequences from USA, Mexico, Western Europe, and Southeast Asia. (https://www.google.com/maps/d/edit?mid=zQfm5yljIn54.k7PW6v6h1pJI&usp=sharing)

**Table 4:** Amino acids substitutions, insertions and deletions in M1 and M2 proteins of RS influenza virus pH1N1.

| Strains | M1 |  |  |  |  |  |  |  |  |  |  | M2 |  |  |  |  |  |  |
| --- | --- | --- | --- | --- | --- | --- | --- | --- | --- | --- | --- | --- | --- | --- | --- | --- | --- | --- |
|  | Substitutions |  |  |  |  |  |  |  | Insertions |  |  | Substitutions |  |  |  |  |  |  |
|  | 47 | 54 | 73 | 91 | 96 | 196 | 197 | 198 | 74/75 | 197/198 | 200/201 | 54 | 77 | 85 | 86 | 91 | 94 | 95 |
| FJ969513 A/California/04/2009 | K | P | G | N | A | S | E | Q | 0 | 0 | 0 | R | Q | D | V | F | I | E |
| KU143699 A/Brazil/RS-1805/2009 | * | * | * | * | * | * | * | * | 0 | 0 | 0 | * | * | * | * | * | * | * |
| KU143701 A/Brazil/RS-1892/2009 | * | * | A | * | * | * | T | T | R | K | S | * | * | * | * | * | * | * |
| KU143695 A/Brazil/RS-2009/2009 | * | * | A | * | * | * | T | T | R | K | S | * | * | * | * | * | * | * |
| KU143697 A/Brazil/RS-2014/2009 | * | * | * | * | * | * | * | * | 0 | 0 | 0 | * | * | * | * | * | * | K |
| KU143694 A/Brazil/RS-2543/2009 | * | * | * | * | * | * | * | * | 0 | 0 | 0 | * | * | * | * | * | * | * |
| KU143691 A/Brazil/RS-2656/2009 | * | * | * | * | T | R | deletion | T | 0 | 0 | S | L | * | * | * | * | * | * |
| KU143689 A/Brazil/RS-2763/2009 | * | * | * | * | * | * | * | * | 0 | 0 | 0 | * | * | * | * | * | * | * |
| KU143703 A/Brazil/RS-2905/2009 | * | * | * | Q | * | * | * | * | 0 | 0 | 0 | * | * | * | * | * | * | * |
| KU143706 A/Brazil/RS-3012/2009 | * | * | * | * | * | * | * | * | 0 | 0 | 0 | * | * | * | * | * | * | * |
| KU143696 A/Brazil/RS-3082/2009 | * | * | * | * | * | * | * | * | 0 | 0 | 0 | * | * | * | * | * | * | * |
| KU143700 A/Brazil/RS-3093/2009 | * | * | * | * | * | * | * | * | 0 | 0 | 0 | * | * | * | * | * | * | * |
| KU143688 A/Brazil/RS-3435/2009 | * | * | * | * | * | * | * | * | 0 | 0 | 0 | * | N | G | C | L | * | * |
| KU143702 A/Brazil/RS-3466/2009 | * | * | * | * | * | * | * | * | 0 | 0 | 0 | * | * | * | * | * | * | * |
| KU143698 A/Brazil/RS-3538/2009 | T | Y | * | * | * | * | * | * | 0 | 0 | 0 | * | * | * | * | * | * | * |
| KU143690 A/Brazil/RS-3908/2009 | * | * | * | * | * | * | * | * | 0 | 0 | 0 | * | * | * | * | * | * | * |
| KU143705 A/Brazil/RS-5081/2009 | * | * | * | * | * | * | * | * | 0 | 0 | 0 | * | * | * | * | * | * | * |
| KU143692 A/Brazil/RS-5377/2009 | * | * | * | * | * | * | * | * | 0 | 0 | 0 | * | * | * | * | * | * | * |
| KU143693 A/Brazil/RS-3335/2009 |  |  |  |  |  |  |  |  |  |  |  | * | * | * | * | * | K | * |
| KU143704 A/Brazil/RS-3504/2009 |  |  |  |  |  |  |  |  |  |  |  | * | * | * | * | * | * | * |

\* No amino acid change.

Seventeen HA sequences obtained from RS samples are available under the GenBank accession numbers MG784976–MG784982 and MG785397–MG785406. These sequences were compared to all sequences available on IRD until July 2016, disregarding the year of influenza virus sequence or the host. The sequences most identical to those obtained in this study were chosen and retrieved to yield a total of 796 sequences. All similarities obtained were for 2009, 2010, and 2011 sequences, and only three were swine-like sequences. Three phylogenetic trees were constructed until obtaining the most similar sequences to yield a total of 258 sequences. The third phylogenetic tree showed only four groups (Figure 5) and was used to construct the interactive map (https://www.google.com/maps/d/u/0/edit?hl=pt-BR&mid=1zn3GhGcRZHYGyct__jIsERPA0ZDQ5Nj4&ll=5.411776974245612%2C0&z=2). The map obtained (Figure 6) showed the worldwide spread (Figure 6A) of HA segment compared to HA sequences of this study and high similarity to other Brazilian (Figure 6B) HA. Distribution of the samples was primarily from the middle to the east of RS (Figure 6C). Only amino acid substitutions were observed in these sequences when compared to California/04/2009 (Table 5), Brazil/RS-3900/2009 Q32E, Brazil/RS-3466/2009 A38E, Brazil/RS-5377/2009 D65E, Brazil/RS-2529/2009 G105E, Brazil/RS-5081/2009 T113K, Brazil/RS-2014/2009, Brazil/RS-2584/2009, and Brazil/RS-3869/2009 Q136H. Brazil/RS-3093/2009 had the most different sequence with N137D, I138M, H139Q, and I141T substitutions. Almost all sequences harbored the substitution I164V, except for Brazil/RS-5377, which harbored a substitution at I167V. The only patient death occurred with Brazil/RS-3222/2009 harboring only one substitution (I164V).

**Figure 5:**
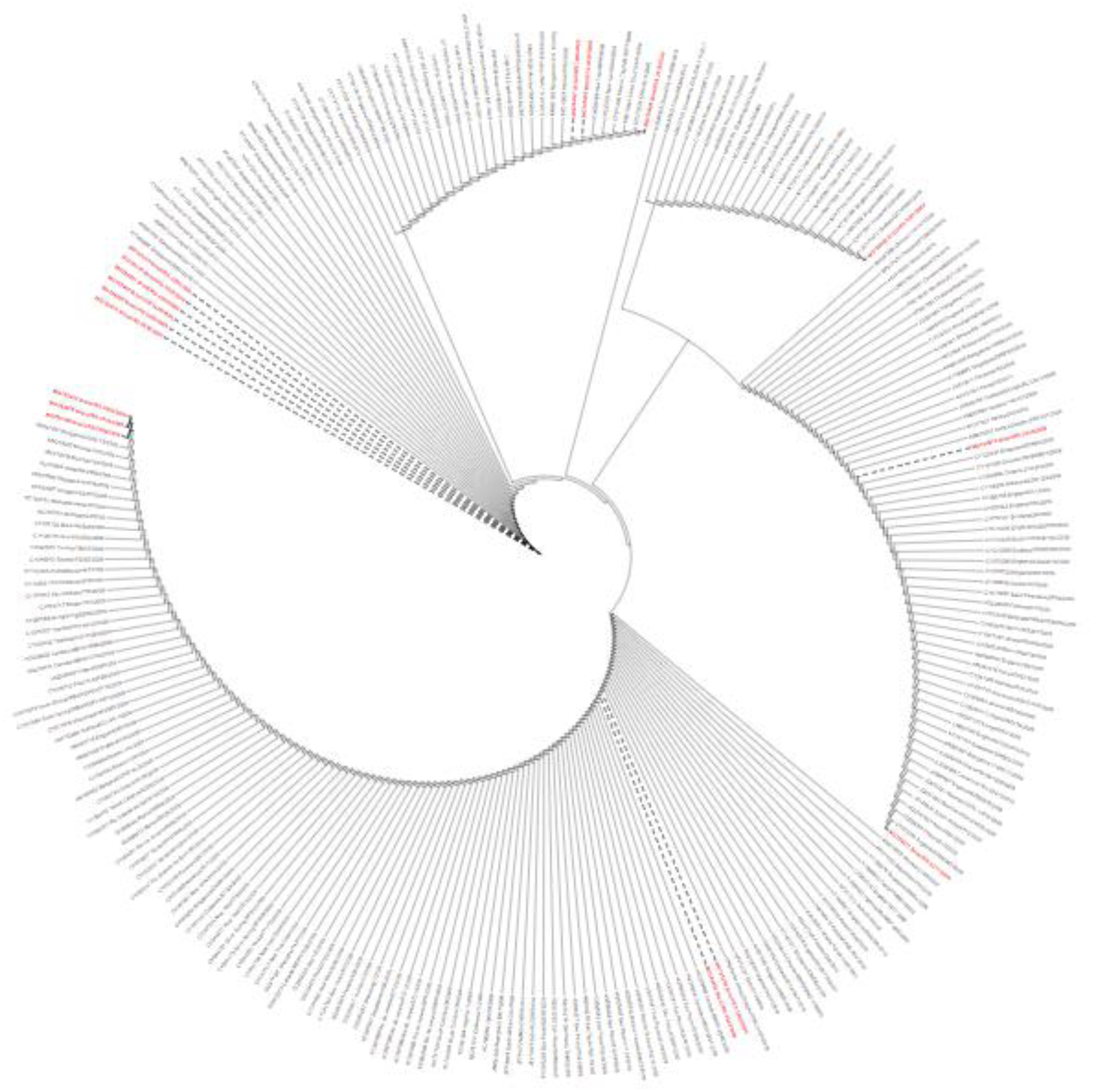
Final phylogenetic tree with 258 of the most similar sequences related to 17 HA RS sequences (red writing). The most divergent sequence is MG784980 showing four mutations which led to the formation of a new clade.

**Figure 6:**
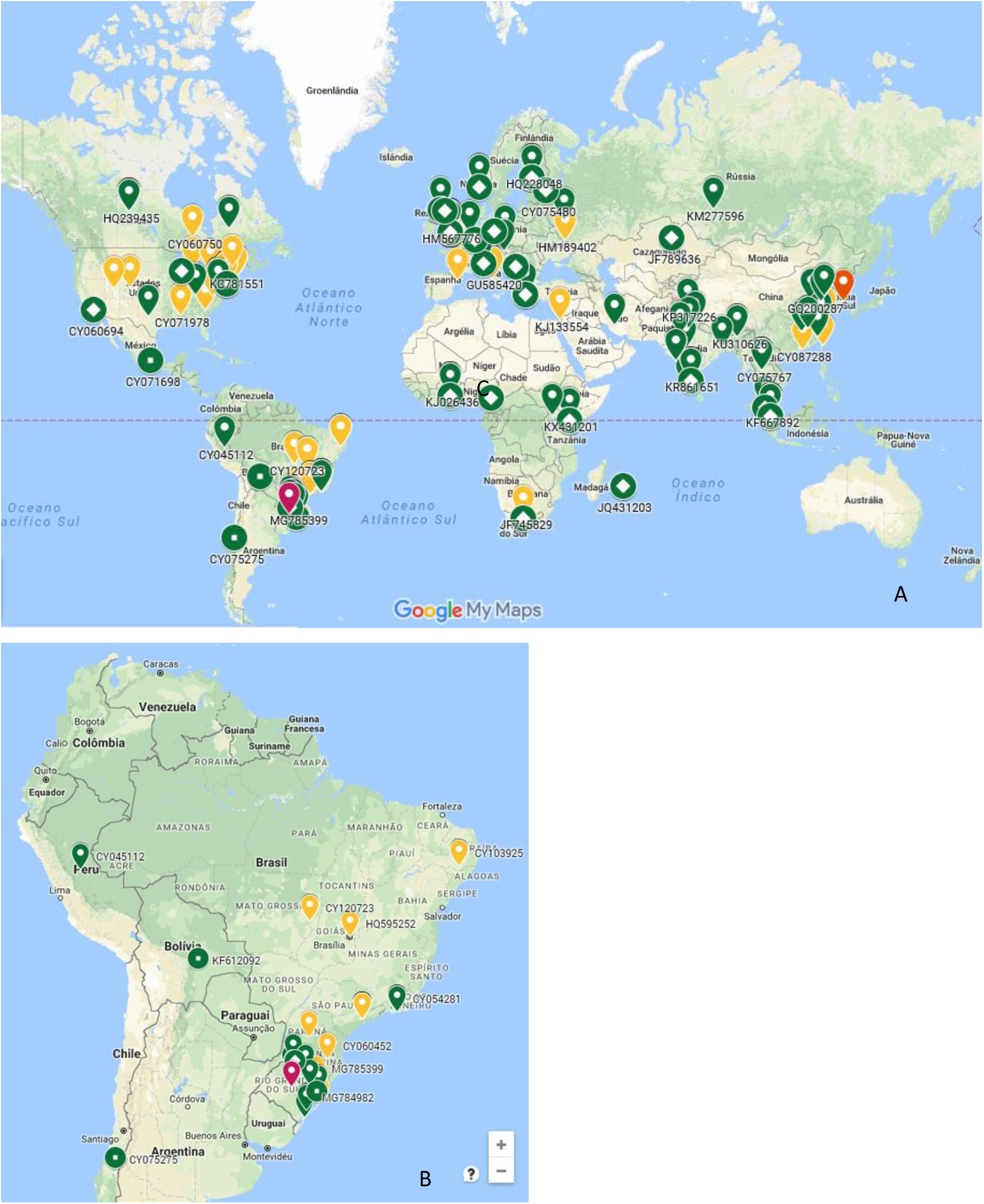
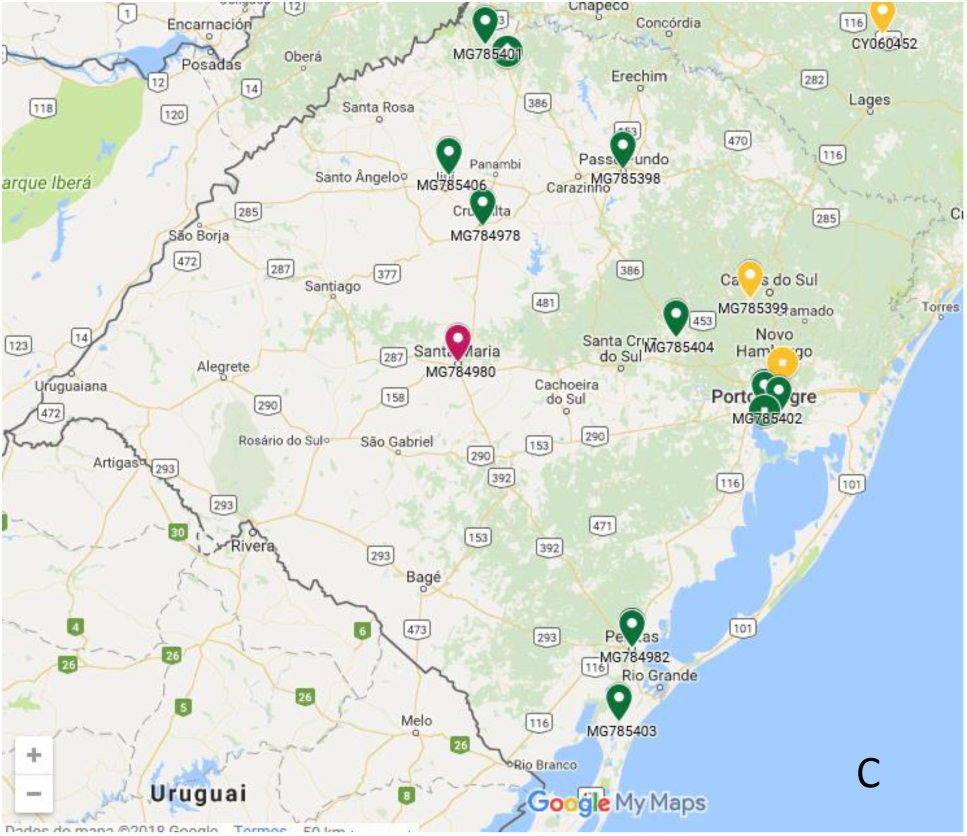
Map of HA most similar sequences obtained from Influenza Research Database to Rio Grande do Sul State (RS). Distribution of HA most similar sequences around the world (A), most similar sequences of Brazil (B) and distribution of samples in RS (C).

**Table 5:**
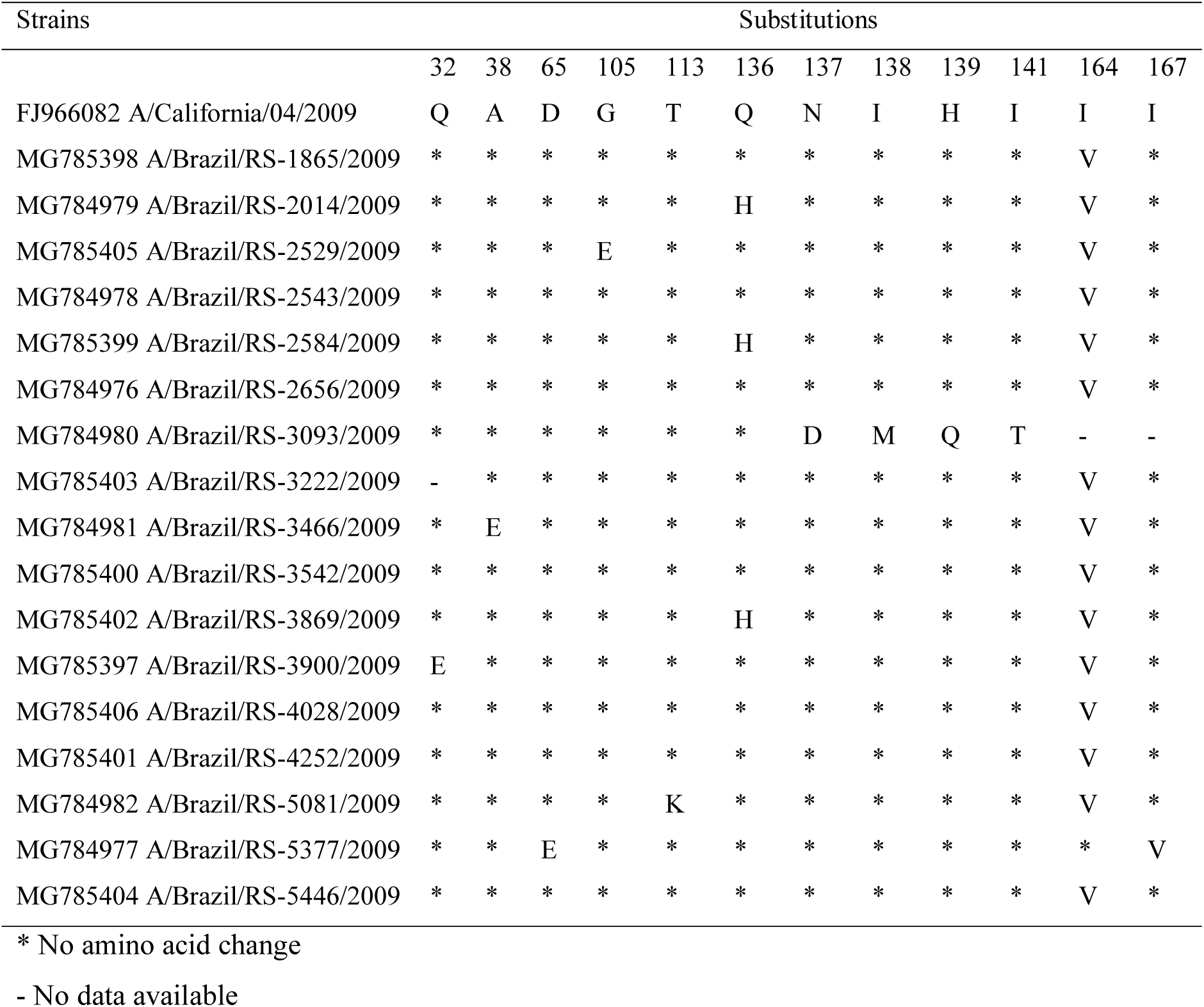
Amino acids substitutions in HA protein of RS influenza virus pH1N1.

### Epidemiology analysis

Although pH1N1 presents no danger to the world, in Brazil, this virus is still monitored by Health Ministry as it exists between circulating viruses and has resulted in higher mortality rates over the years. In 2014, there was an increase in the number of cases of H3N2, overcoming cases of pH1N1 in all regions of Brazil. However, in percentage, the number of deaths was lower than that caused by pH1N1. The virus influenza A H3N2 presented a proportion of deaths of approximately 9% in 2013 and 2014, whereas influenza A pH1N1 virus presented a mortality rate of 20.6% in 2013 and 35% in 2014 (Table 6). The number of cases in southeast and south of Brazil, highlighting the states of South and São Paulo, from 2009 to 2015 are shown in Table 6. It has been observed that Paraná has the highest number of cases in the south, which pertained in 2010. In 2010, RS reported no evidence of pH1N1. However, in 2011, RS presented cases of this virus again, becoming the state with the highest number of cases, unlike other states where the numbers were declining since 2009. In 2012, Santa Catarina, the state just above RS, was the state with the most substantial number of cases, surpassing Sao Paulo in 2013, with a decrease in the number of cases in the three states of south. According to the symptoms observed in RS A(H1N1)pdm09 epidemiological data (Figure 7), chills was the most frequent and more related symptom. Chills showed a direct relationship with fever, a group of symptoms related to each other, viz., arthralgia–myalgia, sore throat–arthralgia, conjunctivitis, and rhinorrhea. The second frequent symptom was rhinorrhea pertaining to chill, conjunctivitis, and cough. There was no significant relationship between metabolic diseases, immunosuppression, heart diseases, and lung disease with other symptoms.

**Figure 7:**
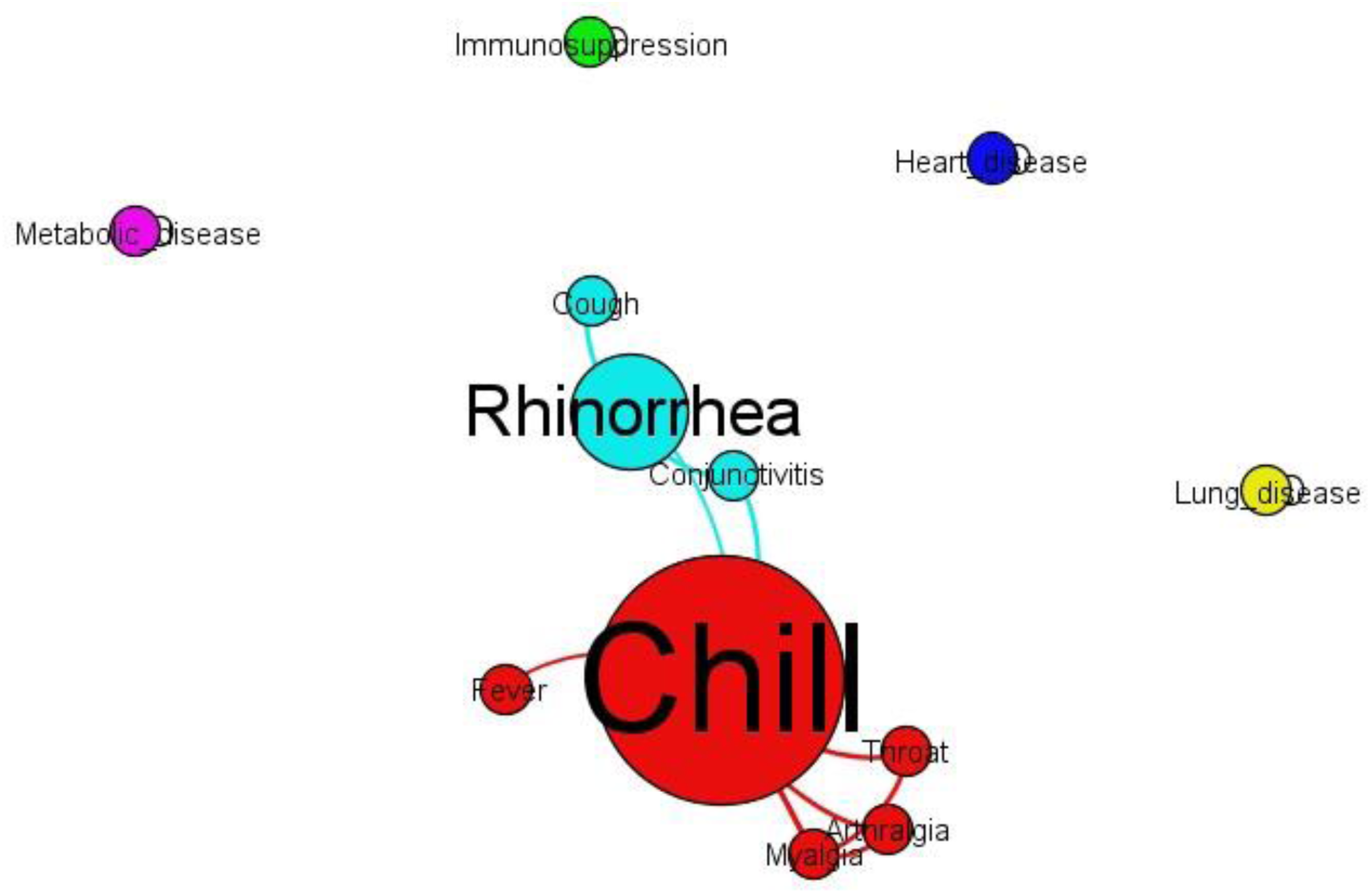
Relationship between symptoms showing chill (biggest red circle) as the most frequent and rhinorrhea (biggest blue circle) as second most frequent and related symptom. Metabolic disease, immunosuppression, heart disease and lung disease didn’t show any relation during A(H1N1)pdm09 in Rio Grande do Sul.

**Table 6:** Cases of influenza A pH1N1 from 2009 to 2014 in Southeast and South of Brazil.

| Brazilian region | State | Year |  |  |  |  |  | 2015 |
| --- | --- | --- | --- | --- | --- | --- | --- | --- |
|  |  | 2009 | 2010 | 2011 | 2012 | 2013* | 2014 |  |
| Southeast | MG | 1810 | 8 | 26 | 134 | 425 | 31 | 5 |
|  | ES | 110 | 1 | 1 | 0 | 17 | 1 | 0 |
|  | RJ | 2777 | 3 | 5 | 4 | 52 | 22 | 0 |
|  | **SP** | **7407** | **108** | **28** | **370** | **1972** | **117** | **14** |
| South | **PR** | **30650** | **344** | **2** | **621** | **353** | **46** | **31** |
|  | **SC** | **2155** | **20** | **7** | **743** | **225** | **21** | **50** |
|  | **RS** | **2592** | **0** | **113** | **520** | **333** | **29** | **0** |

Brazilian States: MG - Minas Gerais, ES - Espírito Santo, RJ - Rio de Janeiro, SP - São Paulo, PR - Paraná, SC - Santa Catarina, RS - Rio Grande do Sul.

## Discussion

The pandemic H1N1 virus (2009) is a reassortment between North American triple- reassortant classical swine virus and Eurasian avian-like swine influenza virus (Morens *et al*. 2009, Zimmer *et al*. 2009), and in this analysis, it was crucial to consider avian and swine influenza viruses independently. Strain California/04/2009 was chosen for comparison because it is a model of transmission (Munster *et al*. 2009) and was also used for annotation.

The small number of amplified samples and an even lower number of overlapping sequences can be explained by the problem of maintenance of samples as discussed by Veiga *et al*. (2012). During the pandemic of 2009, the number of patients throughout the state of RS was considerable and there was no proper way of collecting and storing the samples (Veiga *et al*. 2012).

Samples Brazil/3335/2009 and Brazil/3504/2009 showed the highest similarity to Guangdong/55/2009 (HQ011424). When this sequence was analyzed in IRD, other 12 similar sequences of segment M were found. One of these sequences was KC876544, H3N2, from Argentina. Since RS borders Argentina, many of the cases in the state were from this country, explaining the diversification of this segment in RS. All the RS sequences, similar to HQ011424 and KC876544, showed S31N substitution related to decreased resistance to amantadine (Lan *et al*. 2010). Although the NA segment is important for influenza virus virulence, it is not crucial for virus genome assembly (Gao *et al*. 2008). According to Gao *et al*. (2012), virus lacking the segments PB2, M, or NP exhibit low replication rate and are easily lost during passages in eggs, as these segments seem to recruit other segments during genome packaging. Silent mutations in the M segment can result in packaging defect in all other segments (Hutchinson *et al*. 2008). M1 is an import protein in virus assembly and is responsible for association with influenza virus RNA and ribonucleoprotein (RNP) for controlling nuclear export and import of RNP and some functions in transcription inhibition (Baudin *et al*. 1994, 2001; Bui *et al*. 1996, Huang *et al*. 2001; Whittaker *et al*. 1995). Furthermore, M1/M2 can change the balance of HA and NA, affecting one or both glycoproteins (Campbell *et al*. 2014). Moreover, M from pH1N1 is essential for virion morphology contributing to HA recognition, and increased NA activity improves virus transmission (Campbell *et al*. 2014). Experiments using guinea pigs demonstrated that strains without pH1N1 M could not transmit the virus, whereas those containing NL + M + NA + HA exhibit high transmission even with a virus dose of 100 plaque forming units (PFU) (Campbell *et al*. 2014). According to van Wielink *et al*. (2012), combined mutations in M and NP lead to an average decrease in virus titer, whereas mutations in M itself increase the virus titer, indicating that M alone determines enhanced replication. NS1 is expressed at high levels after infection and facilitates virus replication (Hale *et al*. 2008). One of the goals of NS1 is mRNA splicing of the M segment (Robb and Fodor 2012). The increased titer of delNS1 was due to the M1 amino acid substitution at M86V, a region near to helix six domain, which is a positively charged surface region (van Wielink *et al*. 2012, Sha and Luo 1997). Adaptation of M-segment mutations could compensate for the absence of NS1 protein (delNS1), and these modifications can increase the expression of M1/M2 ratio, even if M2 has silent mutations and M1 with silent or nonsilent mutations that affect M1 mRNA splicing efficiency (van Wielink *et al*. 2012). M1 and M2 proteins can alter NA Vmax through alterations in NA incorporation, distribution, or presentation at the virion surface because the NA cytoplasmic tail interacts in the transmembrane region with M1 protein (Campbell *et al*. 2014, Ali *et al*. 2000, Barman *et al*. 2001, Mitnaul *et al*. 1996).

In the matrix segment, it was found that substitutions in M1 and M2 were significant in virulence. M2 has demonstrated resistance to amantadine and rimantadine in case of S31N substitution (Ilyushina *et al*. 2005), and all RS samples exhibited this trait. This substitution is present in Guangdong/55/2009 and swine/Jangsu/48/2010, and both strains are related to the samples analyzed in this study. Swine/Jangsu/48/2010 was related to Brazil/3012/2009, and Guangdong/55/2009 was nearly related to Brazil/RS-3335/2009 and Brazil/RS-3504/2009 in the first tree construction with all 466 sequences using NJ phylogenetic test and was directly related to Brazil/RS-3504/2009 in the second phylogenetic tree. The patient from whom the sample Brazil/RS-3335/2009 was obtained was the only one who died, indicating highest divergence from all sequences, similar to samples one and two from Beijing. Substitution at 16 and 55 positions of M2 exhibited enhanced transmission in humans. Sample Brazil/RS-2656/2009 showed substitution at position 55, in R54L position, which could also be involved in increased transmission. Fan *et al*. (2009) demonstrated that M gene from H5N1 avian influenza viruses contributes to virulence difference in mice, whereas other genes could attenuate pathogenicity. It is important to consider the geographical position of RS. Presence of an influenza A H3N2 M segment related to pH1N1 M segment from RS is observed because of the representativeness of this virus reassortment in the pandemic of 2009, which is similar to swine/texas4199-2/1998 (H3N2) (Chou *et al*. 2011; Webby *et al*. 2000). The most divergent sequences were obtained in M1 of Brazil/RS-3335/2009 and Brazil/RS-3504/2009. Many differences in sequences were obtained in M1 of Brazil/RS-1892/2009, Brazil/RS-2009/2009, Brazil/RS-2656/2009, and M2 Brazil/RS- 3435/2009. Sequences Brazil/RS-2905 and Brazil/RS-3538/2009 showed one and two substitutions in M1, respectively, and Brazil/RS-2014/2009, Brazil/RS-2656/2009, Brazil/RS-3335/2009 showed one substitution in M2 each. M1 and M2 could have different morphology as some of the samples exhibited signature virion morphology, particularly in the M1 sequence. According to Elleman and Barclay (2004), some of the amino acid morphology determinants in M1 are at residues 41, 95, 218, whereas Bourmakina and Garcia-Sastre (2003) found residues 95 and 204 in M1 as the determinants. Roberts *et al*. (1998) found that M1 and M2 were responsible for virion morphology, including 41 amino acids of M1. Campbell *et al*. (2014) found 13 amino acid differences, including residues 41, 207, 209, 214 in M1, which was similar to model of Elleman and Barclay (2004) but included the difference of 14 amino acids in M2.

The highest similarity in the HA segment was obtained between Brazil/RS-2529/2009 and CY075275 from Chile, Brazil/RS-5081/2009 and JN171873 from Quebec, and Brazil/RS-5377/2009 and CY070245 from England. CY075275 and JN171873 are similar to GQ166223 from China, a sample that showed a high level of replication and mutation of inosine at position 32 (Xu *et al*. 2011). The only sample with a mutation at this site was Brazil/RS-3900/2009 but for glutamic acid. Strains Brazil/RS-2014/2009, Brazil/RS-2584/2009, Brazil/RS-3093/2009 (sequence MG784980), and Brazil/RS- 3869/2009 showed substitutions in the sialic acid ligation site (Al-Maihdi 2007). The most divergent sequence was MG784980, with four mutations in the sialic acid ligation region.

The classical swine influenza virus (SIV) is similar to the 1918 pandemic virus and remained antigenic and genetically conserved in the USA until the introduction of H3N2 in 1998 (Vicent *et al*. 2008). In Europe, an avian-like virus, H1N1, was predominant in swine until the introduction of H3N2 reassortment in the 1980s (VanReeth 2007). Swine have a receptor for both human and avian influenza viruses and, thus, have become an essential player for interspecies transmission. In Brazil, pH1N1 is established in swine populations and may become endemic in the country (Rajão *et al*. 2013). An experiment using California/04/2009 (H1N1) and swine/Texas/4199-2/1998 (H3N2) showed that only viruses harboring the M segment from California/04/2009 exhibited high transmission through the air in guinea pigs (Chou *et al*. 2011). California/04/2009 is highly transmitted through aerosols, whereas swine/Texas/4199-2/1998 did not exhibit this capability. These findings could explain the late substitution of pandemic H1N1 to seasonal H3N2 in RS. Epidemiological data (Brasil 2012a; Brasil 2012b; Brasil 2013; Brasil 2014; Brasil 2015) have probably shown a decrease in cases of pH1N1 in comparison to H3N2. Apparently, the H3N2 virus is more infectious than pH1N1 as it has affected more patients that the pandemic virus, according to this study. This competition could lead to the extinction of the pandemic virus in Brazil. The most common symptoms were chill and rhinorrhea, but not fever, which is usual in case of virus infection. Fever is almost always associated with an immunological response to infection, and this result could indicate why H1N1pdm09 spread worldwide so quickly.

Brazilian SIV HA and NA network suggested a common origin for all virus isolates, irrespective of the region or host (Rajão *et al*. 2013), which is not the same for H1N1pdm2009. In a previous study, Sant’Anna *et al*. (2013) found that most of the Brazilian influenza A pH1N1 sublineages from RS belong to clade 7 (Nelson 2009), assuming that multiple sublineages were introduced in Brazil. M sequences of this study showed different origins and a new clade origin, which was observed in HA as well with Brazil/RS-3093/2009. As M is of avian origin, high divergence in the sequences could be due to migratory birds that allow new recombination of the virus in avian and transmission to humans. RS is also an important place for bird birth and migration during summer and winter. According to Morrison and Ross (1989), there are representatives of species of birds such as *Calidris sp., Pluvialis sp., Tryngites sp., Sterna sp., Cygnus sp., Dendrocygna sp., Gallinula sp., Plegladis sp., Netta sp*., and *Coscoroba sp*. It is possible that the M segment originated from Eurasian avian-like swine virus and underwent natural selection or even, as discussed by Sant’Anna *et al*. (2013), founder effect. Thus, as proposed by Ozaki *et al*. (2014), specific host cell phenotypes may differentially influence virus replication because reassortments between influenza A H6N1 and influenza A H6N2 PB2 and M repressed replication in the chicken trachea. In another study, avian mutations in the M segment increased virulence in mice as M1 protein contributes to the virulence of H5N1 avian influenza viruses (Fan et al. 2009).

The HA sequences (classical swine segment) showed less divergent sequences than M sequences (avian-like segment). Notably, IRD had 19,018 M and 34,565 HA sequences until January 16, 2018 in its system. According to this data, the number of HA sequences is almost the double of M sequences in the genomic database, and this difference could explain the similarities of HA sequences, which was not observed in M sequences.

Epidemiological data along with molecular analysis of samples obtained from the 2009 pandemic may explain the increased number of cases in RS from 2011 to 2012 and the consequent increase in cases in 2012 in the states of Santa Catarina and Paraná, culminating in the highest number of cases in Sao Paulo in 2013. São Paulo is the most populous state in the country and shows some winter characteristics observed in the south, such as change in temperature, which in addition to air pollution could explain the high number of cases in the state. It is possible that differentiated viruses grouped in a clade apart from the phylogenetic tree have somehow prevailed in the state and have mutated to a more virulent form since 2011. As shown by Rajão *et al*. (2017), there are many reassortments between H1N1pdm09 and H3N2 in swine. These reassortments include at least one segment of H1N1pdm09 in H3N2, particularly the M segment, and HA is stable, as shown in this study for humans. As discussed above, RS, just like Santa Catarina, is a bird migration area, which may have influenced the maintenance of this virus in the country along with new mutations in the M segment as the M segment originates from avian or swine. Santa Catarina has most of the swine breeding sites of the country and could have led to the reassortments between avian and swine viruses.

## Acknowledgments

We thank CAPES for the financial support.

